# The HLA-E/NKG2A axis identifies a therapeutic vulnerability in BCG-unresponsive bladder cancer

**DOI:** 10.1101/2024.09.02.610816

**Authors:** B. Salomé, H. Yu, D. Ranti, Y.A. Wang, C. Bieber, P-E Gabriel, I. Duquesne, T. Strandgaard, B. Hug, Sean Houghton, H. Ravichandran, E. Merritt, S. Bang, A. Demetriou, Z. Li, S. V. Lindskrog, A. Rangel, D.F. Ruan, J. Daza, J. Cris Ingles, R. Rai, R. Fernandez-Rodriguez, S. Izadmehr, G. Doherty, A. Narasimhan, A.M. Farkas, M. Tran, J. Park, J. Qi, M. Patel, D. Geanon, G. Kelly, R. Zhang, R.M.de Real, B. Lee, K. Nie, S. Miake-Iye, K. Angeliadis, E. Radkevich, A. Soto, R.P. Sebra, S. Smith, M. Skobe, E. Clancy-Thompson, D. Palmer, S. Hammond, B. D. Hopkins, P. Wiklund, J.J. Bravo-Cordero, R. Brody, Z. Chen, S. Kim-Schulze, L. Dyrskjøt, O. Elemento, A. Tocheva, W-M. Song, R. Mehrazin, N. Bhardwaj, M.D. Galsky, J.P. Sfakianos, A. Horowitz

**Author notes:** **Corresponding Authors:** Amir Horowitz, PhD, Associate Professor of Immunology & Immunotherapy and Oncological Sciences Lipschultz Precision Immunology Institute and Tisch Cancer Institute Icahn School of Medicine at Mount Sinai 1425 Madison Ave., Box 1630. New York, New York 10029, |, John P. Sfakianos, MD Associate Professor of Urology, Icahn School of Medicine at Mount Sinai 1425 Madison Ave., L6-58, Box 1272 New York, New York 10029. These authors contributed equally (B. Salomé, H. Yu, D. Ranti). Co-Senior author (J.P. Sfakianos, A. Horowitz).

## Abstract

**Background:** Bacillus Calmette-Guérin (BCG) is the standard of care treatment for high-risk non-muscle-invasive bladder cancer (NMIBC), yet many patients develop recurrent disease despite evidence of ongoing immune activation. We investigated mechanisms of immune escape in BCG-unresponsive tumors and evaluated the therapeutic potential of targeting the HLA-E/NKG2A axis.

**Methods:** Single-cell RNA sequencing, spatial immunophenotyping, proteomic profiling, and functional ex vivo assays were performed using tumors and urine samples from patients with BCG-naïve and BCG-unresponsive NMIBC.

**Results:** BCG-unresponsive tumors were enriched for HLA-E-expressing malignant cells compared with BCG-naïve tumors. Increased HLA-E expression was associated with enhanced IFN-γ signaling and was induced by IFN-γ stimulation in primary tumor cells and bladder cancer tumor lines. Spatial analyses demonstrated accumulation of NKG2A^+^ NK and CD8 T cells in proximity to HLA-E^high^ tumor cells, with increased NKG2A:HLA-E interactions in BCG-unresponsive tumors. Despite high expression of cytotoxic mediators, NKG2A^+^ effector cells displayed impaired degranulation. Blockade of NKG2A with monalizumab restored degranulation of and cytotoxicity by tumor-infiltrating lymphocytes in autologous tumor co-cultures.

**Conclusions:** BCG-unresponsive NMIBC tumors are enriched for HLA-E-expressing tumor cells and NKG2A^+^ effector lymphocytes, with increased engagement of the HLA-E/NKG2A axis within the tumor microenvironment. These findings identify the HLA-E/NKG2A axis as a therapeutic vulnerability and provide a rationale for clinical evaluation of NKG2A blockade as a bladder-sparing strategy for patients with BCG-unresponsive disease.

## INTRODUCTION

In patients with high-risk non–muscle-invasive bladder cancer (NMIBC), Bacillus Calmette-Guérin (BCG) remains the standard first-line bladder-preserving immunotherapy^1^. Upon transurethral resection of the bladder tumor (TURBT), BCG is administered as a six-week induction course and induces a broad inflammatory response characterized by recruitment and activation of innate and adaptive immune cells, including monocytes, neutrophils, dendritic cells, T cells, and NK cells^2,3^. Despite its use for over 40 years, up to 40% of patients develop recurrent disease, and up to 13% progress to muscle-invasive disease^4,5^. Radical cystectomy remains the standard curative option for BCG-unresponsive patients, but is associated with substantial morbidity and a significant impact on quality of life^6,7^.

The biological mechanisms that enable tumor persistence despite BCG-induced immune activation remain poorly understood. While PD-L1 upregulation is observed in a subset of BCG-unresponsive tumors^8^, clinical responses to PD-1/PD-L1 blockade in NMIBC have been modest and PD-L1 expression alone does not consistently predict treatment outcomes^5,8–11^. These observations suggest that additional immune-evasion pathways contribute to therapeutic resistance within the bladder tumor microenvironment (TME).

HLA-E is a non-classical MHC class I molecule that suppresses cytotoxic lymphocyte activity through engagement of the inhibitory receptor NKG2A expressed on NK cells and subsets of CD8 T cells^12–16^. Blockade of NKG2A enhances anti-tumor immunity in preclinical models^13,16^ and has demonstrated clinical activity in combination with PD-L1 blockade in patients with advanced solid tumors, such as non-small cell lung cancer^17,18^. We previously identified the HLA-E/NKG2A axis as a key regulator of NK-like CD8 T cell function in bladder cancer, supporting a broader role for this pathway in bladder tumor immune surveillance and immune escape^15^.

Here, we investigated the role of the HLA-E/NKG2A axis in BCG-unresponsive NMIBC. Using single-cell transcriptomics, spatial immunophenotyping, proteomic profiling and functional *ex vivo* analyses, we demonstrate that BCG-unresponsive tumors are enriched for HLA-E expressing malignant cells and accumulate NKG2A⁺ NK and CD8 T cells in close proximity to HLA-E^high^ tumor regions. We further show that IFN-γ signaling is associated with HLA-E expression and BCG recurrence, and that IFN-γ induces HLA-E expression in primary tumors and bladder cancer cell lines. Finally, NKG2A blockade restores cytotoxic effector-cell function in patient-derived samples, identifying the HLA-E/NKG2A axis as a therapeutic vulnerability in BCG-unresponsive bladder cancer.

## RESULTS

### HLA-E is enriched in BCG-unresponsive tumors

We previously demonstrated that HLA-E expression in bladder tumors decreases as the disease progresses, although expression levels were highly variable at the non-muscle invasive (NMI) stage^15^. Here, we investigated the impact of BCG therapy on HLA-E expression in NMIBC tumors. We first profiled bladder tumor cell composition by single-cell RNA sequencing (scRNAseq) in treatment-naive NMI bladder tumors (n=3) and in bladder tumors from patients who recurred after BCG therapy (n=4) (Table S1). UMAP clustering analysis revealed seven major subsets of tumor cells (n=18,520 cells) (Figure 1A, Table S2) with varying *HLA-E* expression and distribution depending on BCG exposure (Figure 1B). Cluster B7 displayed the highest expression of *HLA-E* and was enriched in BCG-unresponsive tumors.

**Figure 1.**
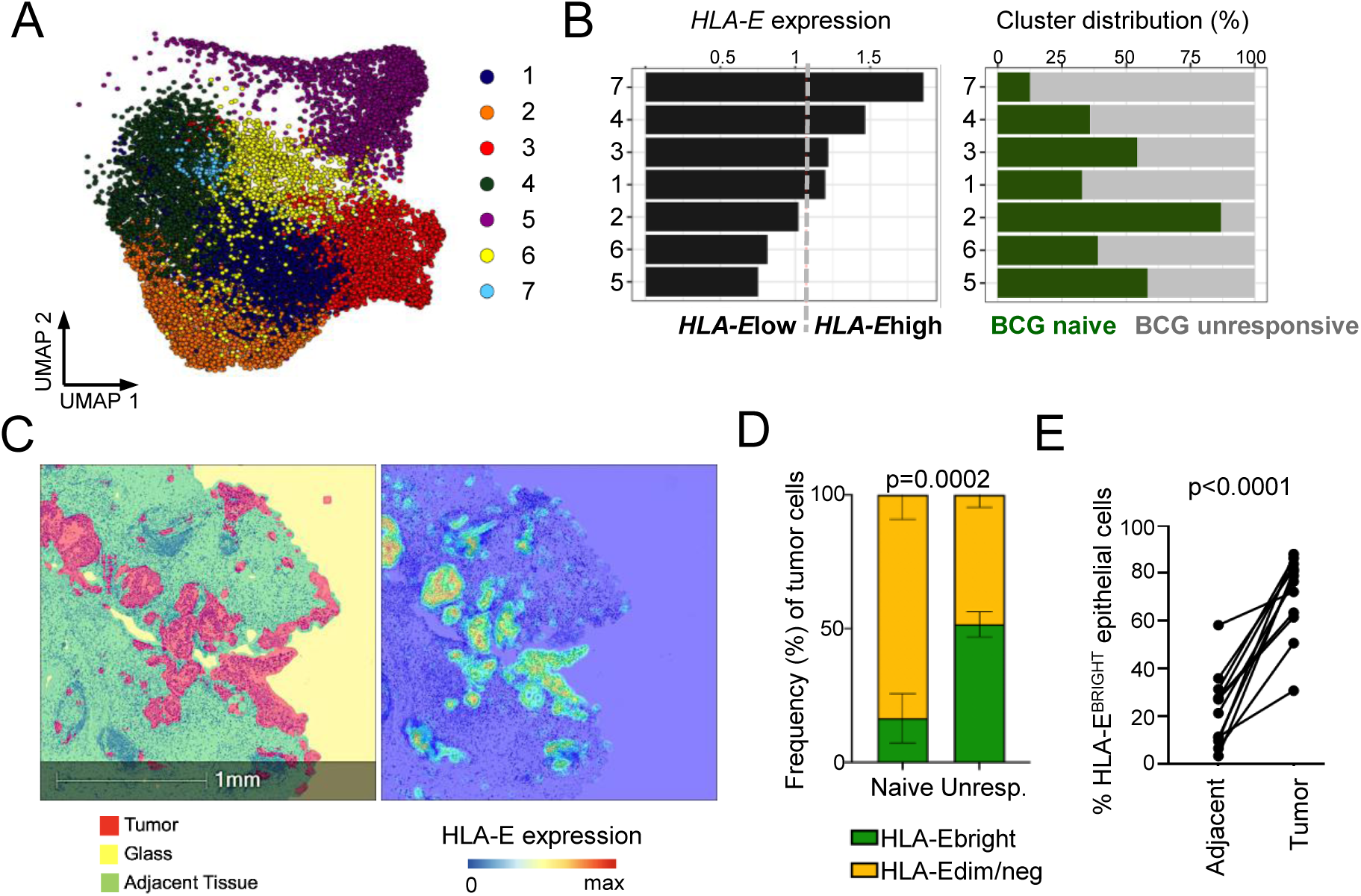
HLA-E is enriched in BCG-unresponsive tumors (A-B) Single-cell RNA sequencing was performed on ex vivo bladder tumors (n=3 BCG-naïve, n=4 BCG-unresponsive) and tumor cells selected for further analyses. **(A)** UMAP visualization of bladder tumor cells from unsupervised clustering. Each color represents one cluster. **(B)** Average expression of HLA-E per cluster (left) and distribution of the bladder tumor cell clusters in BCG-naïve and BCG-unresponsive NMIBC tumors (right). **(C-E)** IHC was performed on bladder tumor and adjacent non-involved tissues from BCG-naïve and BCG-unresponsive NMIBC patients. **(C)** Representative digital pathology analyses identifying tumor, adjacent tissue along with exposed areas of glass to be excluded from subsequent analyses (left) and gradient of HLA-E tumor expression (right) **(D)** Summary analysis of frequency of tumor cells that are HLA-E-bright and HLA-E-dim/negative in BCG-naïve (n=17) and BCG-unresponsive (“unresp.”, n=24) NMIBC tumors. The p-value was obtained using an independent two-sided t-test. **(E)** Summary analysis of frequency of HLA-E^bright^ epithelial cells in tumor and adjacent, non-involved bladder tissues from BCG-unresponsive patients with NMIBC. The p-value was obtained using a paired t-test. Lines show matching samples from a same donor.

We then profiled HLA-E expression at the protein-level by multiplexed immunohistochemistry (IHC) on 41 bladder tumor sections (840,348 cells in total) from BCG-naïve (n=17) and BCG-treated (n=24) patients. We identified tumor cells and adjacent non-tumor epithelial cells as positive and negative for pan-cytokeratin, respectively (Figure 1C). HLA-E^bright^ tumor cells were significantly more abundant in BCG-treated unresponsive tumors than in BCG-naïve tumors (43.6% vs 10.3%, p=0.0002; Figure 1D). Within BCG-unresponsive tumors, HLA-E expression was significantly higher in tumor cells than in adjacent, non-involved urothelium (71.2% vs 21.6% HLA-E^bright^ epithelial cells, p<0.0001; Figure 1E).

### NKG2A^+^ effector cells accumulate near HLA-E^bright^ tumor cells in BCG-unresponsive tumors

We next evaluated the abundance and spatial distribution of NKG2A+ effector cells in NMIBC tumors. Multiplex immunohistochemistry identified NKG2A expression on both CD3^+^ (T cells) and CD3^-^ (NK cells) and enabled assessment of their proximity to HLA-E^bright^ and HLA-E^dim/neg^ tumor cells (Figure 2A). NKG2A+ cells were significantly enriched in BCG-unresponsive tumors compared with BCG-naïve tumors (Figure 2B). In both treatment-naïve and BCG-unresponsive tumors, NKG2A was represented in similar proportions in NK cells and T cells. (Figure 2C).

**Figure 2.**
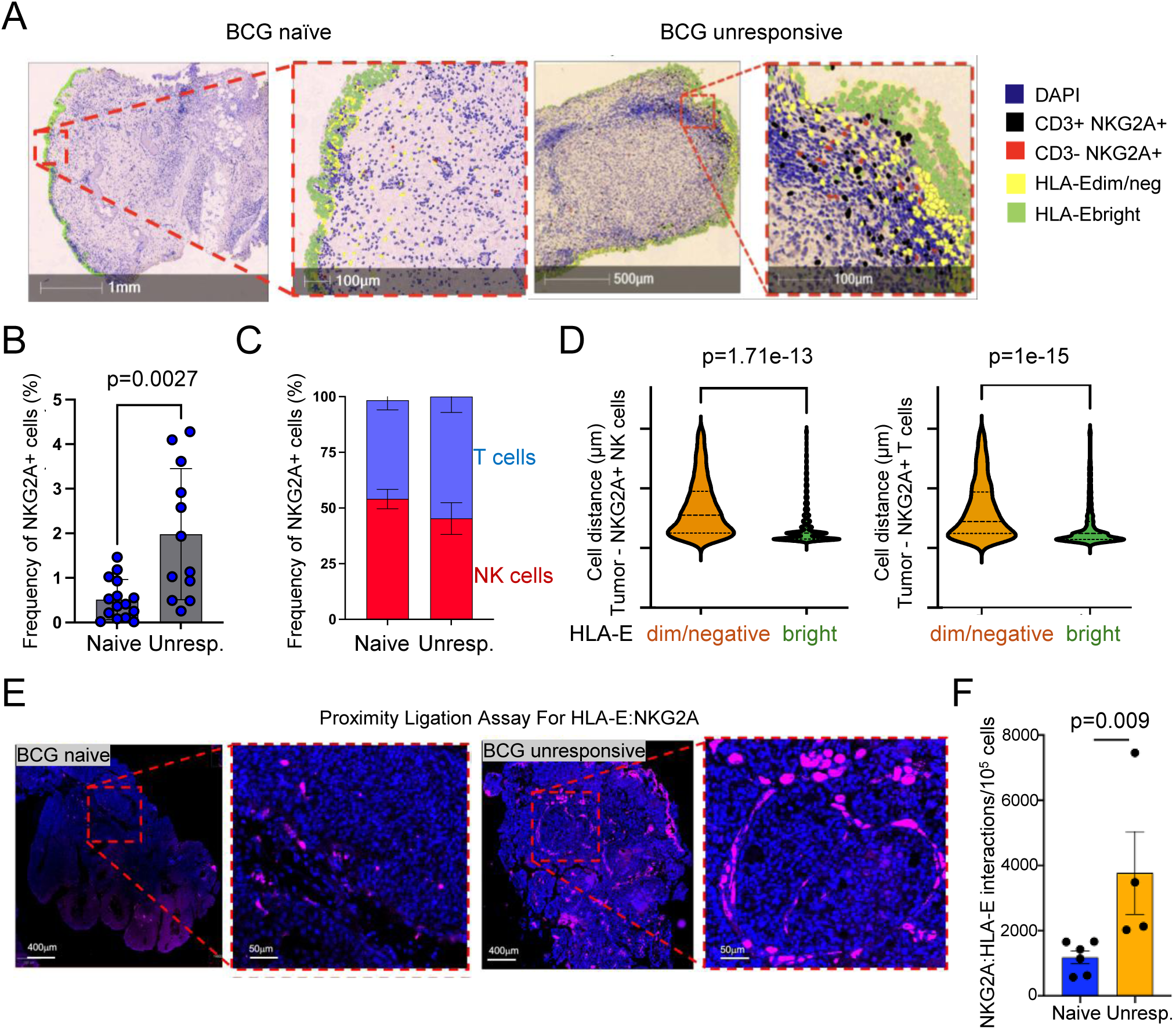
NKG2A+ effector cells accumulate near HLA-Ebright tumor cells in BCG-unresponsive tumors (A-D) IHC was performed on NMIBC tumors (n=41 tumors). **(A)** Representative digital pathology analysis on one BCG-naïve and one BCG-unresponsive NMIBC tumor section highlighting nuclear expression of DAPI and identification of CD3^+^NKG2A^+^ T cells, CD3-NKG2A^+^ NK cells and tumor cells with bright or dim/negative expression of HLA-E. **(B)** Frequencies of NKG2A^+^ cells in BCG-naïve (n=17) and BCG-unresponsive (“unresp.”, n=24) tumors. **(C)** Frequencies of NK cells within the NKG2A^+^ cell compartment in BCG-naïve (n=17) and BCG-unresponsive (n=24) tumors. **(D)** Proximity analysis measuring the cell distance from CD3^-^ NKG2A^+^ NK cells or CD3^+^NKG2A^+^ T cells and tumor cells, depending on the tumor cell expression of HLA-E. P-values were assessed via independent two-sided t-test. **(E-F)** Interactions between HLA-E and NKG2A were profiled using an immunofluorescence-based proximity ligation assay in bladder tumors from NMIBC patients (n=10). **(E)** Representative staining in one BCG-naïve (left) and one BCG-unresponsive (right) patient. **(F)** Summary analysis of the interactions between HLA-E and NKG2A in BCG-naïve (n=6) and BCG-unresponsive (n=4) tumors. The p-value was assessed via independent two-sided t-test.

Spatial analyses demonstrated that both NKG2A⁺ NK cells and NKG2A^+^ T cells preferentially localized near HLA-E^bright^ tumor cells compared with HLA-E^dim/neg^ tumor cells (NK cells: p=1.7×10^-13^, T cells: p=1×10^-15^; Figure 2D). To determine whether these spatial relationships reflected engagement of the HLA-E/NKG2A axis, we performed proximity ligation assays on tumor sections from BCG-naïve (n=6) and BCG-unresponsive (n=4) patients (Figure 2E). HLA-E and NKG2A interactions were significantly increased in BCG-unresponsive tumors (p=0.009; Figure 2F). Together, these findings demonstrate enrichment of NKG2A^+^ effector cells and increased engagement with HLA-E in BCG-unresponsive NMIBC.

### IFN-ɣ signaling is associated with HLA-E expression and BCG-unresponsive disease

We next investigated transcriptional programs associated with HLA-E expression in bladder tumor cells. Differential expression analysis of *HLA-E*^high^ and *HLA-E*^low^ tumor cells identified enrichment of inflammatory and interferon-response pathways, including IFN-ɣ signaling, in *HLA-E*^high^ cells (Figure 3A).

**Figure 3.**
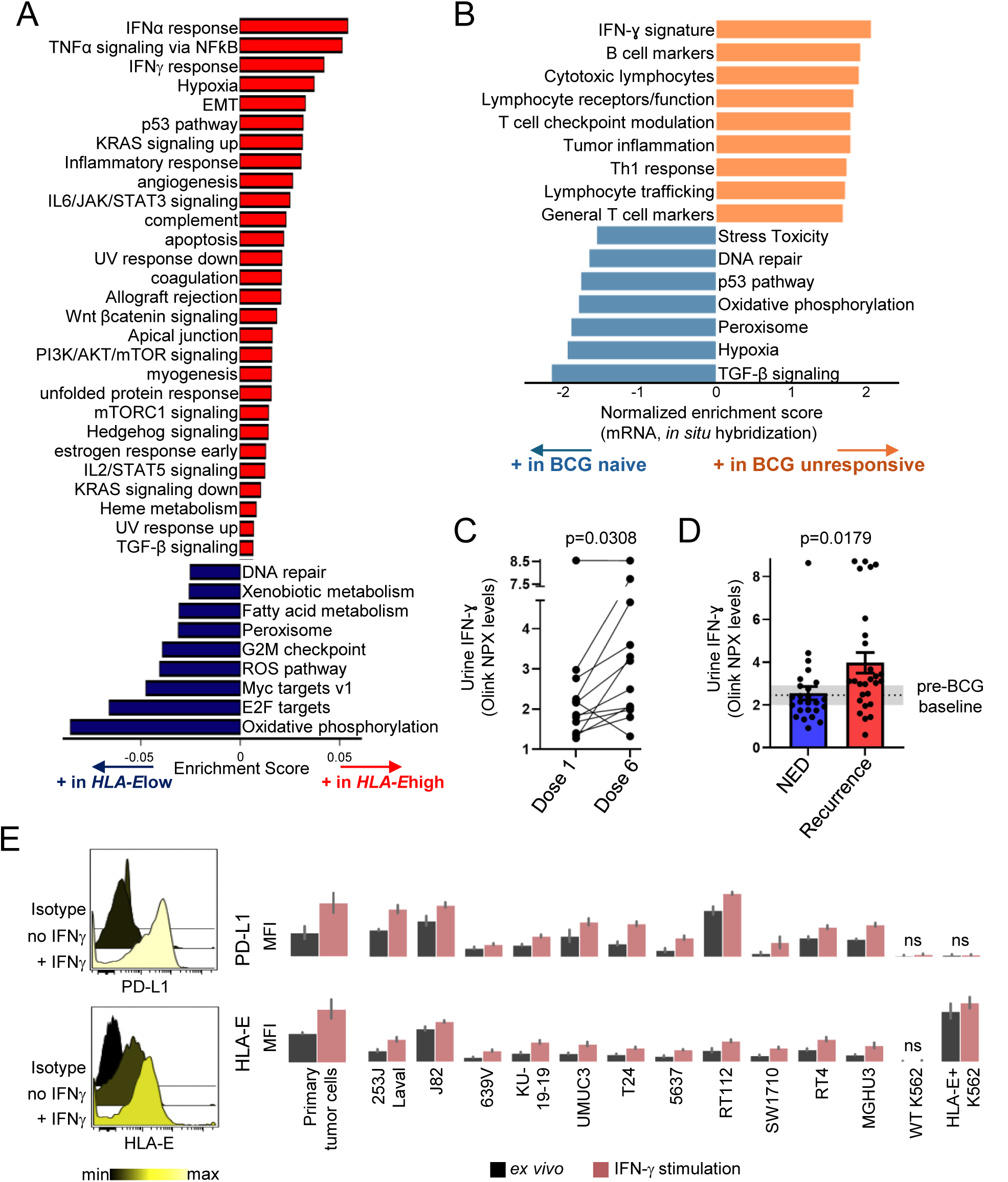
IFN-ɣ drives HLA-E expression and associates with BCG resistance. **(A)** Single-cell RNA sequencing was performed on ex vivo bladder tumors (n=3 BCG-naïve, n=4 BCG-unresponsive) and tumor cells selected for further analyses. Pathway analysis of Hallmark gene networks that are significantly differentially expressed on HLA-E^high^ and HLA-E^low^ bladder tumor cells. **(B)** Targeted mRNA gene-set enrichment analysis was performed using High-Throughput Genomics (HTG). All gene sets represented were statistically enriched (p<0.05) in BCG-naïve (n=20) or BCG-unresponsive tumors (n=20), as assessed with an independent two-sided t-test. **(C-D)** Olink protein analysis was performed on urine supernatants of patients undergoing BCG therapy. **(C)** IFN-ɣ concentrations were compared between dose 1 and dose 6 of the first induction cycle and **(D)** between non-evidence of disease (absence of tumor) and recurrence cases of BCG-treated patients (right panel). Lines show matching samples from a same patient. P-values were obtained with paired (C) or unpaired (D) t-tests. **(E)** Flow cytometry was performed on primary CD45^-^ tumor cells from NMIBC patients and immortalized bladder tumor lines *ex vivo* and upon 24 hour stimulation with rhIFN-ɣ. Representative (left) and summary (right) expression of PD-L1 and HLA-E in triplicate experiments.

To determine whether these findings extended to BCG-unresponsive disease, we performed targeted transcriptomic profiling of BCG-naive (n=17) and BCG-unresponsive (n=19) tumors (Table S1). BCG-unresponsive tumors were enriched for inflammatory pathways, including IFN-γ response, tumor inflammation, lymphocyte trafficking and modulation of T cell functions (Figure 3B, Table S2).

Given the enrichment of IFN-γ signaling in both *HLA-E*^high^ tumor cells and BCG-unresponsive tumors, we examined the relationship between IFN-γ and HLA-E expression. Urinary proteomic analyses demonstrated a significant increase in IFN-γ during the six-dose BCG induction cycle, which was validated in an independent cohort (Figure 3C, S). IFN-γ concentrations were also higher in patients who subsequently developed recurrence than in patients with no evidence of disease following BCG treatment (Figure 3D).

To assess the effect of IFN-γ on tumor cells, primary NMIBC tumor cells (n=10) and bladder tumor lines were cultured with recombinant IFN-γ. Wild-type (“WT”) K562 cells, lacking class I HLA, remained unresponsive, while HLA-E-transduced K562 cells served as positive controls. Across primary tumors and bladder tumor lines, IFN-γ consistently increased HLA-E expression (p<0.05) (Figure 3E). As expected, IFN-γ also induced PD-L1 expression on primary tumor cells and all bladder tumor lines (p<0.05). Together, these findings identify IFN-γ as a potential driver of HLA-E upregulation in BCG-unresponsive NMIBC.

### NKG2A blockade restores cytotoxic effector functions in BCG-unresponsive NMIBC

To determine whether NKG2A^+^ effector cells represent a therapeutically targetable population in BCG-unresponsive disease, we characterized their cytolytic potential by mass cytometry of immune cells from urine and primary tumor samples (n=8: 5 urine and 3 tumor samples). NKG2A expression was associated with increased expression of cytotoxic mediators, including Granzyme A and Perforin in both NK and CD8 T cells, and Granzyme B in NK cells (Figure 4A-B).

**Figure 4.**
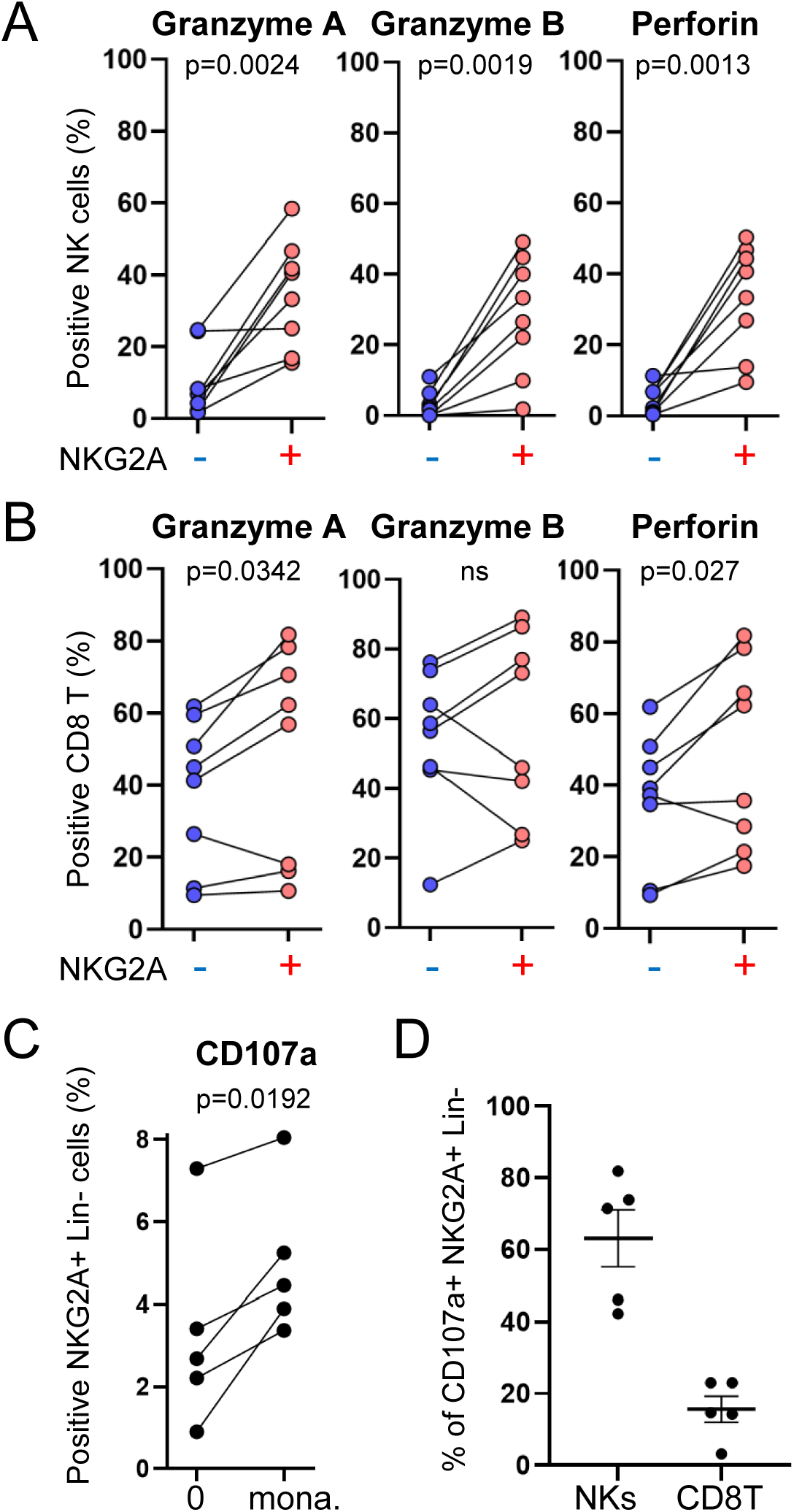
NKG2A blockade restores anti-tumor cytotoxicity in BCG-unresponsive patients (A-B) Mass cytometry was performed on *ex vivo* cells from n=8 bladder tumors or urine in BCG-unresponsive patients. **(A)** Granzyme A, Granzyme B and Perforin expression in NKG2A+/- NK cells and **(B)** NKG2A**+/-** CD8 T cells. **(C-D)** CD45+ cells from n=5 BCG-unresponsive bladder tumors or urine were co-cultured for 5h at a 1:1 ratio with matching CD45- tumor cells, in the presence of brefeldin, monensin and CD107a, before proceeding with mass cytometry staining. **(C)** Frequencies of degranulating (CD107a+) cells within the live CD45+ NKG2A+ Lineage- cell compartment, in the presence or absence of monalizumab (anti-NKG2A). **(D)** Frequencies of NK cells and CD8 T cells within the degranulating live CD45+ NKG2A+ Lineage-cell compartment.

We next assessed the functional consequences of NKG2A blockade using co-cultures of tumor-infiltrating immune cells with autologous tumor cells from BCG-unresponsive patients (n=5). Addition of the anti-NKG2A antibody, monalizumab, significantly increased degranulation of tumor-infiltrating NKG2A^+^ immune cells (n=5) (2.3-fold increase, p=0.03) (Figure 4C). Phenotypic analysis revealed that degranulating NKG2A^+^ cells consisted predominantly of NK cells (67%), with a smaller contribution from CD8 T cells (16%) (Figure 4D). These findings identify NKG2A^+^ effector cells as a functionally constrained cytotoxic population in BCG-unresponsive NMIBC and demonstrate that NKG2A blockade enhances their effector function.

## DISCUSSION

BCG remains the cornerstone bladder-preserving therapy for high-risk NMIBC, yet a substantial proportion of patients ultimately develop recurrent tumors despite evidence of ongoing immune activation. The mechanisms underlying immune escape following BCG therapy remain incompletely understood. In this study, we identify the HLA-E/NKG2A axis as a clinically relevant mechanism of immune evasion in BCG-unresponsive NMIBC. Using complementary single-cell, spatial, proteomic and functional approaches, we demonstrate that recurrent tumors are enriched for HLA-E-expressing malignant cells, accumulate NKG2A^+^ NK and CD8 T cells, and display increased NKG2A:HLA-E interactions. Importantly, blockade of NKG2A restores effector-cell functions in patient-derived samples, identifying this pathway as a viable therapeutic target in BCG-unresponsive disease.

A central observation of our study is that recurrent, BCG-unresponsive tumors remain highly inflamed despite failure of immune-mediated tumor control. *HLA-E*^high^ tumor cells were enriched for interferon-responsive transcriptional programs, while BCG-unresponsive tumors displayed increased inflammatory signaling, including IFN-ɣ pathway activation. Consistent with these findings, urinary IFN-ɣ increased during BCG induction and remained elevated in patients who subsequently developed recurrence. Furthermore, exogenous IFN-ɣ directly induced HLA-E expression in primary NMIBC tumors and bladder tumor lines.

These findings should be interpreted within the broader context of IFN-ɣ biology. In a meta-analysis spanning 18 tumor types, including bladder cancer, inflammatory mediators such as IFN-ɣ were associated with expression of inhibitory immune checkpoints, including PD-L1/L2^19,20^. Despite emerging evidence supporting pro-tumorigenic roles for IFN-ɣ, its anti-tumor functions remain essential for effective responses to immune checkpoint blockade. Anti-tumor inflammation therefore exists along a continuum in which sustained immune activation can simultaneously promote tumor control and induce compensatory inhibitory pathways. Together, our findings support a model in which persistent inflammatory signaling following BCG therapy promotes HLA-E expression within recurrent tumors. These findings suggest that more is not necessarily better for every patient and provide a rationale for future studies evaluating whether biomarkers of inflammatory activation could guide personalization of the current six-week BCG induction regimen.

HLA-E is a non-classical MHC class I molecule that suppresses cytotoxic lymphocyte activity through engagement of the inhibitory receptor NKG2A. We further demonstrate that NKG2A^+^ effector cells represent a therapeutically relevant population in BCG-unresponsive NMIBC. NKG2A^+^ NK and CD8 T cells were enriched in recurrent tumors, preferentially localized near HLA-E^high^ tumor cells, and displayed increased interactions with HLA-E. NKG2A expression identified functionally competent NK and CD8 T cells with elevated expression of cytotoxic mediators, including granzymes and perforin. Importantly, blockade of NKG2A with monalizumab significantly increased degranulation in autologous tumor co-cultures. Together, these findings indicate that NKG2A^+^ effector cells remain amenable to functional reinvigoration and provide a mechanistic rationale for therapeutic targeting of this pathway.

The translational relevance of these findings is supported by the clinical development of NKG2A blockade. Monalizumab, a humanized anti-NKG2A antibody, has demonstrated biological and clinical activity in combination with immune checkpoint blockade in solid tumors. In the Phase II COAST trial, addition of monalizumab to durvalumab improved progression-free survival compared with durvalumab alone in patient with unresectable stage III NSCLC^17^. Although subsequent studies have yielded mixed results, these findings established NKG2A as a clinically actionable checkpoint target. Our data extend these observations to BCG-unresponsive NMIBC and suggest that therapeutic strategies targeting NKG2A may complement existing checkpoint blockade approaches by simultaneously enhancing NK- and CD8 T-cell-mediated anti-tumor immunity.

Building upon the findings presented here, we recently initiated the Phase II ENHANCE trial to evaluate durvalumab and monalizumab in patients with high-grade BCG-unresponsive (or BCG-exposed) NMIBC (clinicaltrials.gov ID: NCT06503614). In addition to assessing clinical activity, this study will evaluate baseline tumor HLA-E expression and NKG2A abundance as candidate biomarkers for future patient selection and therapeutic response.

This study has several limitations. Although it represents one of the largest molecular analyses of BCG-treated NMIBC reported to date, sample sizes remain modest for several experimental cohorts. In addition, because bladder tissue is typically sampled following BCG therapy only in patients with evidence of recurrence on cystoscopic evaluation, longitudinal comparison of matched pre- and post-treatment tumors between responders and BCG-unresponsive patients is limited.

In conclusion, our findings identify the HLA-E/NKG2A axis as a mechanism of immune escape in BCG-unresponsive NMIBC. The enrichment of HLA-E^high^ tumor cells, accumulation of NKG2A^+^ cytotoxic effector cells and restoration of effector function following NKG2A blockade provide a strong rationale for clinical evaluation of NKG2A-targeted therapies as bladder-preserving treatment strategies for patients with BCG-unresponsive disease.

## ACKNOWLEDGMENTS

We thank Mary Anne O’Donnell (Precision Immunology Institute, Icahn School of Medicine at Mount Sinai) for critically reviewing the manuscript and Deepta Bhattacharya’ (University of Arizona) for kindly providing HLA-E+ K562 tumors. Additionally, we thank Emily Mace and Everardo Solloa-Hegewisch (Columbia University) for expert guidance on imaging analyses. We acknowledge the expertise and assistance of the Dean’s Flow Cytometry Core at Mount Sinai (RRID:SCR 027701). The A.H. and J.P.S labs were supported by funding from P30CA196521, R01CA269954, R21CA274148, BCAN (Bladder Cancer Advocacy Network) No. 961726, R01CA301050, and R21AI130760A. The N.B. lab was supported by funding from the Department of Defense Peer Reviewed Cancer Research program Translational team Award No. W81XWH1910269 and from the Parker Institute for Cancer Immunotherapy No.AGR-11611SOW1. The J.J.B-C lab was supported by funding from R01CA244780, R61CA278402, the Irma T. Hirschl Trust and the Emerging Leader Award from the Mark Foundation. The L.D lab was supported by funding from the Danish Cancer Society. The O.E lab was supported by the following grants: NCI250233, OT2CA297580, U24CA264032, R01AI187598, R01CA271915, U01DA058527, R01CA271619, R01CA271545, R01NS111997, R01NS127984, R01CA269954, U19AI168632, OT2OD032720, 80NSSC23K0832, CGCATF2023100030, HHWF240412, and LLS702723. A.N.M.R.S was part of the A.H lab as part of an existing collaboration with Dr. Leonardo Oliveira Reis (UroScience, University of Campinas-UNICAMP, Campinas, São Paulo, Brazil) and National Council for Scientific and Technological Development (CNPq-Brazil Edital 26/2021- Process 402562/2022-4) and received a CNPq-PDE fellowship (Process 201230/2024-0). Authors affiliated with the Tisch Cancer Institute were supported by the National Cancer Institute Cancer Center Support Grant P30CA196521.

## AUTHOR CONTRIBUTIONS

B.S., H.Y., D.R., Y.A.W., C.B., J.D., N.B., M.G., J.P.S., and A.H. conceived the project and experiments, analyzed the data and wrote the manuscript. J.P.S., R.M., P.W., M.G., and R.B. provided access to the human samples. B.S., H.Y., D.R., Y.A.W, C.B, P-E.G., I.D., T.S., B.H., S.H., E.M., S.B., A.D., Z.L., S.V.L., A.R., D.F.R., J.D., J.C.I., R.R., K.A., E.R., H.R., S.H., R.F-R., S.I., G.D., A.N., A.M.F., M.T., J.P., A.S., R.P.S., J.J.B-C., O.E performed the experiments or analyzed data. J.Q., M.P., D.G., G.K., R.Z., R.M.D.R., B.L., S.K-S, K.N., S.M-I., acquired sample data using Olink proteomics or mass cytometry. S.S., M.S., E.C-T., D.P., S.H., B.D.H., Z.C., L.D., A.T., W-M.S. provided intellectual input.

## METHODS

### RESOURCE AVAILABILITY

#### Lead contact

Further information and requests for resources and reagents should be directed to and will be fulfilled by the Lead Contact Amir Horowitz.

#### Materials availability

This study did not generate new unique reagents.

#### Data and Code availability

The single-cell RNAsequencing data generated by the authors have been uploaded to the Gene Expression Omnibus (GSE276014) and will be made publicly available upon publication of this manuscript.

The algorithms used in this study will be made available at https://github.com.

### EXPERIMENTAL MODEL AND SUBJECT DETAILS

#### Human subsets

Patients at Mount Sinai Hospital (MSH) were enrolled in the study following Institutional Review Board (IRB) approval (protocol 11-01565). 11-01565 covers the use of patient tissues in a biorepository and allows for prospective collection of blood, urine, and tissue samples from enrolled patients. Formalin-fixed paraffin-embedded (FFPE) blocks from BCG patients were obtained retrospectively from the biorepository and prospectively for patients receiving treatment. For prospective patients, samples were collected on the day of surgery and throughout BCG immunotherapy. Tumor samples were taken at every possible timepoint. BCG naïve was defined as any patient who had yet to receive BCG, regardless of past treatment with other chemotherapies. BCG-unresponsive was defined as any patient with recurrent tumors following at least five of six induction doses of BCG at time of first evaluation.

#### Cell lines

HLA-E+ and Wild-type K562 cell lines were kindly provided by Deepta Bhattacharya and propagated as recently described^21^. HLA-E+ K562 cells were generated using the AAVS1-EF1a donor plasmid containing the coding sequence for human HLA-E. The K562 were electroporated using a Bio-Rad Gene Pulse electroporation system. HLA-E+ cells were sorted to >98% purity. Bladder cancer cell lines were provided by John Sfakianos: 253J(RRID: CVCL_7935), 639V(RRID:CVCL_1048), 5637(RRID: CVCL_0126), J82(RRID: CVCL_0359), KU-19-19(RRID: CVCL_1344), MGHU3(RRID: CVCL_9827), RT4(RRID: CVCL_0036), RT112(RRID: CVCL_1670), SW1710(RRID:CVCL_1721), T24(RRID: CVCL_0554), UMUC3(RRID: CVCL_1783).

### METHOD DETAILS

#### Sample processing

Urine samples from bladder cancer patients were spun down within 30min-1 hour post-collection. The cell-free supernatant was stored at −80C and later used for proteomics analyses. The cell pellet was washed with PBS, filtered through a 40μM cell strainer and resuspended in RPMI-1640 medium supplemented with 10% Fetal Bovine Serum (FBS), Penicillin/Streptomycin (1%) and Glutamax (1%) (“R10 medium”). Tumor tissues obtained from transurethral resections of bladder tumor (TURBT) and cystectomies were placed into RPMI-1640 medium. Tumor tissues were then digested using tumor dissociation enzymes (Miltenyi, 130-095-929) and a GentleMACS machine (program 37C_h_TDK_2) at 37C, filtered through a 40μM cell strainer and resuspended in R10 medium.

#### Multiplex Immunohistochemistry

Sections of tumors for immunohistochemical (IHC) staining were taken at a thickness of 3-mm from formalin fixed paraffin-embedded (FFPE) blocks. H&E-stained sections were performed every 5 - 10 slices. The Ventana Discovery Ultra (Roche Diagnostics) machine was used to automatically bake, deparaffinize, and condition the slices. The RUO Discovery Universal (v21.00.0019) was used to perform chromagen IHC on sequential slices. Primary antibodies included CD3, HLA-E, NKp46, and NKG2A and were utilized for staining on NMIBC tumors. All slices followed the same protocol, which included a 60 minute incubation at 37°C; secondary antibodies using OmniMap HRP or NP DISCOVERY (Roche Diagnostics); signal detection using Discovery OmniMap. Nuclear counterstaining with Mayer’s hematoxylin; and conversion to high-resolution images via the NanoZoomer S10 Digital slide scanner (Hamamatsu).

#### Tumor and CD3^+^ T cell expression and proximity analyses

Prior to analysis, all slides were reviewed and regions of interest were annotated by a board-certified pathologist (R.B.). Tissue artifacts, including torn, folded, and damaged tissue, were excluded from any analyses. The HALO^TM^ (Indica Labs, Inc.) digital image analysis platform, a semi-automated platform using machine learning to segment and label stained sections, was utilized for quantitative analyses. Halo AI^TM^ and train-by-example classification, segmentation, and random forest classification was used to separate chromogenic stains and generate tabular data for downstream analysis. Slide features of each tumor, including cell lineages (tumor, stroma, and immune) and slide features (such as glass) were characterized. Glass was excluded from all downstream analyese. Classified cell classes were tabulated, and positive staining cells were stratified into expression tertiles (dim, moderate, and bright). Calibration for intensity expression was performed using tonsil tissues from healthy human tonsil. In addition to cell counts, total surface area (mm^2^) was recorded to facilitate density calculations. Statistical analyses were performed using Python 3.8.1.

#### Proximity Ligation Assay: sample preparation

The NaveniFlex^TM^ Proximity Ligation Assay (PLA) was performed according to the manufacturer’s instructions using NaveniFlex Tissue MR ATTO647N (Navinci, Sweden). PLA was performed on sections of tumor taken at a thickness of 3-mm from formalin fixed paraffin-embedded (FFPE) blocks. H&E-stained reviewed with pathologist. Briefly, after deparaffinization, rehydration, and antigen retrieval, slides were blocked with Block NT blocking solution (Navinci, NT.1.100.01) for 60 min at 37 °C in a preheated humidity chamber and then incubated with mouse anti-HLA-E (clone: MEM-E/02, Abcam, 1:200) and rabbit anti-NKG2A (clone: EPR23737-127, Abcam, 1:2000) diluted in Diluent 1 NT solution (Navinci, NB.1.100.02) overnight at 4°C. As negative controls, two (tonsillectomy) slides were incubated in antibody diluent with only one primary antibody each. After washing, the slides were incubated with the PLA probes corresponding to the primary antibodies using anti-mouse Navenibody M1 NT (Navinci, NB.1.100.06) and anti-rabbit Navenibody R2 NT (NB.1.100.07) in Diluent 2 NT solution (Navinci, Navinci, NF.1.100.03) for 60 min at 37 °C. Slides were then processed for ligation using reaction 1 reagent containing Buffer 1 NT (Navinci, NB.2.100.17) and Enzyme 1 NT (Navinci, NF.2.100.11) and subsequently reaction 2 reagent containing Buffer 2 NT (NT.2.100.01) and Enzyme 2 NT (Navinci, NF.2.100.15) and incubated for 30 min at 37 °C and 90 min at 37 °C, respectively. The slides were washed and incubated with post-block NT reagent (Navinci, NF.1.100.01) in post-block supplement NT (Navinci, NT.2.100.04) for 30 min at 37 °C, then processed for detection, counter-stained with DAPI, and mounted with coverslips using Prolong Gold Antifade reagent (Invitrogen, P36930).

#### Proximity Ligation Assay: images capture

Images were captured at the Microscopy and Advanced Bioimaging Core of the Icahn School of Medicine at Mount Sinai. A Leica DMi8 (Leica Microsystems, Germany) was equipped with a HC PL APO CS 10x/0.4 (Part Number 506285; Leica Microsystems, Germany) objective lens. A SpectraX fluorescence illuminator (Lumencor, Oregon, USA) with multiple narrow-band light emitting diodes provided illumination (LEDs used: 395/25nm for DAPI, 470/24nm for autofluorescence channel, and 640/30nm for Navinci signal). The microscope and light source were controlled by LAS X software, version 3.7.5.24914 (Leica Microsystems, Germany). For fluorescence excitation, the following illumination settings were used: a 395nm LED set to 50% (147mW at the SpectraX output port) for DAPI signal, captured at 20 milliseconds of exposure; a 470nm LED set to 44% (86mW), captured at 70 milliseconds for an autofluorescence channel; and a 640nm LED set to 100% (231mW), captured at 150 milliseconds for the target signal. A multi-band pass filter set (Part Number 11525366; Leica Microsystems, Germany) was used to separate fluorophore emission (Dichroic 415/490/570/660nm; Emission bands: 430/35, 515/40, 595/40, 720/100nm). Images were captured using a Leica DFC9000GT monochrome camera set to 12-bit depth, 2x2 (4-pixel) binning and “Low Noise” Gain mode. Images were captured in montage at 10% overlap, merged (“Smooth” blending option) and then saved in the proprietary LIF (“Leica Image File”) format before being converted to IMS (Imaris) format for analysis.

#### Proximity Ligation Assay: images analyses

Image analysis was performed using Imaris software 10.1.1 (Oxford Instruments, Concord MA). A surface for the green background channel was created using the surface creation wizard with the following parameters – Enable Region Of Interest = false, Enable Region Growing = false, Enable Tracking = false, Enable Classify = false, Enable Shortest Distance = false, Enable Smooth = true, Surface Grain Size = 2.00 µm, Enable Eliminate Background = false, Active Threshold = true, Enable Automatic Threshold = false, Manual Threshold Value = 1900, Active Threshold B = false. Masked channels were created by subtracting the intensities within the green surface from the blue and far-red channels: the mask intensity was set to 0 for inside the green surface while the outside was set to the original channel’s value. A new surface was created using the surface creation wizard for the masked far-red channel using the following parameters – Enable Region Of Interest = false, Enable Region Growing = false, Enable Tracking = false, Enable Classify = false, Enable Shortest Distance = false, Enable Smooth = true, Surface Grain Size = 2.00 µm, Enable Eliminate Background = false, Active Threshold = true, Enable Automatic Threshold = false, Manual Threshold Value = 1600, Active Threshold B = false. An area filter was applied to this far-red surface to remove surfaces whose area was larger than 50um^2^. A DAPI surface was created using surface creation wizard with the parameters – Enable Region Of Interest = false, Enable Region Growing = true, Enable Tracking = false, Enable Classify = false, Enable Shortest Distance = true, Enable Smooth = true, Surface Grain Size = 2.00 µm, Enable Eliminate Background = true, Diameter Of Largest Sphere = 7.50 µm, Active Threshold = true, Enable Automatic Threshold = false, Manual Threshold Value = 10, Active Threshold B = false, Region Growing Estimated Diameter = 6.00 µm, Region Growing Morphological Split = false, Filter Seed Points = “Quality” above 60.0, Filter Surfaces = “Number of Voxels Img=1” between 10.0 and 500, to obtain individual nulcei within the field of views. Finally, a fourth surface was created by applying the filter – Overlapped Area to Surfaces (Minimum = 0.050 um^2^, Maximum = false) to obtain the DAPI surfaces that were in contact with the far-red channel. The counts of total number of nuclei and nuclei overlapping with the masked far-red surfaces were extracted from the statistics tab of Imaris.

#### Protein concentration measurement

OLINK Proteomics®- inflammation panel and immuno-oncology 92 assays panel were used to profile cell-free urine supernatant from the Mount Sinai and the Aarhus university cohorts, respectively. Cell-free urine supernatant and serum samples were randomized in a 96-well plate. and incubated overnight alongside negative controls with an incubation mix (incubation solution, incubation stabilizer, A-probes, and B-probes) at 4°C. Samples were then incubated with an extension mix (High purity water, PEA solution, PEA enzyme, PCR polymerase) for 5 min and placed on a thermal cycler. Following the thermal cycler, samples were incubated with a detection mix (detection solution, High purity water, detection enzyme, PCR polymerase) and transferred to a chip. Primers were loaded onto the chip, and the chip was profiled using the Fluidigm IFC controller HX with the Fluidigm Biomark Reader. Data were normalized using extension and interplate controls and a correction factor. The resulting data were reported in normalized protein expression (NPX) units on a log2 scale. In order to determine the suitable statistical test, a Shapiro-Wilk’s test was used to assess for normality, and a Kruskal-Wallis test was used in every instance in which one or both samples were not normally distributed. An independent T-test was used in the event both samples were normally distributed. All statistically significant p values were then used to assess adjusted p values via the Benjamini-Hochberg correction, with an alpha of 0.05. All statistically significant genes between the BCG naïve and sixth induction dose time points are shown.

#### IFN-γ stimulation of tumor cells

Cell lines and CD45- isolated primary tumor cells were incubated in media optimized for high viability for 72 hours (RPMI-1640 supplemented with 20% fetal bovine serum). Tumor cells were expanded until they were confluent in two T175 flasks. Following expansion, cells were cultured in 100 ng / mL of IFN-γ for a total of 24 hours in a 24 well plate. Following co-culture, cell lines were trypsinized and primary tumors were gently removed from the solid phase by a cell scraper. HLA-E and PD-L1 levels were assessed via Flow Cytometry. Cells were stained in 4C FACS buffer (phosphate-buffered saline (PBS) with 2% heat-inactivated FBS and EDTA 2 mM) for 30 minutes. Subsequently, cells were washed in PBS, incubated for 20 minutes in a viability dye, washed again with PBS, and resuspended in 2% paraformaldehyde. The experiment was performed in triplicate, with three readouts per cell line per experimental condition. Samples were acquired with an LRS-Fortessa (BD Biosciences), and data were analyzed using the CytoBank software. When staining for HLA-E, cells were first stained 20 minutes with HLA-E prior to staining with additional PD-L1. In CytoBank, several gates were applied to generate the final dataset. A live/dead gate was applied, followed by a gate to remove doublets and isolate singlets. Lastly, the data was arcsinh transformed prior to analysis.

#### Single-cell RNA sequencing: Data preprocessing

Single-cell RNA sequencing (scRNA-seq) analysis was performed using Python and R. After loading, genes expressed in fewer than three cells were excluded from later analyses. Cells with < 200 or > 8000 unique genes, as well as cells containing >20% mitochondrial gene transcripts, were discarded. Doublet cells were screened by scrublet, where expected doublet rate was set at 5.0%, and detected doublet rate was 0.0%. Subsequent data was preprocessed with the Scanpy package and normalization was performed by dividing feature counts for each cell by total counts for that cell, scaling by a factor of 10,000, and natural log transformation. Next, scvi model was set up to correct batch-effects. Then we performed scaling and principal component analysis (PCA) on the batch-corrected data. Using the first 50 principal components (PCs), graph-based clustering and UMAP dimensionality reduction was performed to reveal 16 cell clusters. All scRNAseq analyses were performed using distinct samples without repeated measurements.

#### Single-cell RNA sequencing: Subclustering analyses

We performed subclustering analysis on 18,520 bladder tumor cells using Seurat v5 workflow. We integrated tumor cell transcriptomes across samples using Canonical Correlation Analysis (CCA) integration with the IntegrateLayers() function (dimensional reduction for correction = pca). Leiden clustering (resolution = 0.4) was applied to the shared nearest neighbors (SNN) (dims = 1:10). This resulted in the identification of 9 heterogeneous tumor subclusters, distinctly separated on the UMAP plot. Each subcluster was profiled by the expression of tumor marker genes (EPCAM, UPK2) and cytokines (CXCL1, CXCL2, CXCL3, IFNGR1, etc.). Subcluster B1 was removed from further analysis as it was identified as normal bladder cells with low EPCAM (tumor-related marker genes), and subcluster B8 was removed due to high PTPRC expression, indicating a high presence of immune cells (CD45+). Differentially expressed gene (DEG) analysis (MAST, R version 1.2.1)^22^ and enrichment pathway analysis^23^ further characterized the subclusters, highlighting their distinct biological profiles.

#### Co-cultures with autologous tumor cells

CD45^+^ and CD45^-^ cells were isolated from freshly dissociated bladder tumor cells (CD45^+^, CD45^-^) or from urine cells (CD45^+^) using the EasySep Release Human CD45 positive selection kit (Stemcell, cat#100-0105). CD45+ cells were stained for 15min at room temperature with monalizumab (20ng/mL; gift from AstraZeneca), or resuspended in medium. CD45+ cells were then co-cultured with autologous CD45- cells at a 1:1 ratio during 5 hours in the presence of brefeldin A (BioLegend, 420601), monensin (BioLegend, 420701) and CD107a 147Sm (H4A3 clone, Maxpar) in 96-well plates, 37C, 5% CO2.

#### Mass cytometry antibody preparation

When available, conjugated antibodies were purchased from Fluidigm (CD16 209Bi clone 3G8, CD45 89Y HI30, CD107a 147Sm H4A3). Remaining antibodies were purchased carrier-free and were conjugated in-house using the Maxpar X8 and MCP9 labeling kit (Fluidigm), as per the manufacturer instructions: CD4 142Nd (RPA-T4 clone), CD8 151Eu (RPA-T8), CD56 158Gd (REA196), NKG2A 160Gd (REA110), CD3 166Er (UCHT1), Granzyme A 112Cd (CB9), Granzyme B 171Yb (QA16A02), Perforin 172Yb (REA1061), as well as lineage markers in the 115In channel: CD14 (M5E2), CD19 (HIB19), CD20 (2H7), ɣδTCR (B1).

#### Mass cytometry staining

Cells were incubated during 20 min at 37C in R10 cell medium in the presence of IdU (Maxpar, cat#201127) and Rh103 (Maxpar, cat#201103A). Cells were centrifuged and washed with the Maxpar cell staining buffer (NC0501752) prior to a 3-min incubation on ice with an Fc-blocking reagent (BioLegend, cat# 422302). Samples were then washed and stained with extracellular antibodies for 30 min on ice in cell staining buffer. Samples were then washed and resuspended in Fixation/Perm buffer (Invitrogen, cat#00-5523-00) for 30 min on ice. Cells were then centrifuged, washed with Permeabilization buffer (Invitrogen, cat#00-5523-00) and stained with intracellular antibodies in permeabilization buffer in the presence of Heparin 100U/mL for 30min on ice. Stained cells were washed with permeabilization buffer and resuspended in PBS in the\ presence of PFA 2.4%, saponin 0.08% and Ir 0.125uM (Fluidigm, cat#201192A). Stained samples were finally washed and resuspended in cell staining buffer and data acquired within a week.

#### Mass cytometry sample acquisition and processing

Immediately prior to acquisition, samples were washed with Cell Staining Buffer and Cell Acquisition Solution (Fluidigm) and resuspended in Cell Acquisition Solution at a concentration of 1 million cells per mL containing a 1:20 dilution of EQ normalization beads. The samples were acquired on the Fluidigm Helios mass cytometer using the wide bore injector configuration at an acquisition speed of <400 cells per second. The resulting FCS files were normalized and concatenated using Fluidigm’s CyTOF software. The FCS files were further cleaned using the Human Immune Monitoring Center at Mt. Sinai’s internal processing pipeline. The pipeline removes aberrant acquisition time-windows of 3 s where the cell sampling event rate goes above or below 2 standard deviations from the mean acquisition speed. EQ normalization beads are removed as well as low DNA intensity events. Samples were demultiplexed by calculating the cosine similarity of every cell’s Palladium barcoding channel to every possible barcode used in a batch. Once the cell has been assigned to a sample barcode, a signal-to-noise metric is calculated by taking the difference between its highest and second highest similarity scores. Any cells with low signal-to-noise are flagged as multiplets and removed from that sample. Finally, acquisition multiplets are removed based on the Gaussian parameters Residual and Offset acquired by the Helios mass cytometer.

#### Mass cytometry data analysis

Cytobank was used to assess the frequencies of cells expressing markers of interest. CD8 T cells were defined as live CD45+ Lineage- CD3+ CD4- CD8+, and NK cells as live CD45+ Lineage- CD3- CD56+ cells.

#### Statistical analyses

Statistical analyses were performed using GraphPad Prism 10, R and Python. All tests were two-sided unless otherwise specified. Comparisons between two groups were performed using Student’s t-test or paired t-test as indicated in the figure legends. For transcriptomic analyses, p-values were adjusted using the Benjamini-Hochberg method. Statistical significance was defined as p<0.05.

---

**Table S1.** Clinical cohort characteristics.

| HTG | scRNAseq | Olink | CyTOF | Age at collection | Sample Type | BCG Timeline | BCG response | Days since last BCG |
| --- | --- | --- | --- | --- | --- | --- | --- | --- |
| x |  |  |  | 90 | tumor | Post | resistant | 50 |
| x |  |  |  | 89 | tumor | Pre | resistant | NA |
| x |  |  |  | 81 | tumor | Pre | resistant | NA |
| x |  |  |  | 79 | tumor | Post | resistant | 470 |
| x |  |  |  | 64 | tumor | Pre | resistant | NA |
| x |  |  |  | 64 | tumor | Post | resistant | 48 |
| x |  |  |  | 77 | tumor | Post | resistant | 861 |
| x |  |  |  | 86 | tumor | Post | resistant | 56 |
| x |  |  |  | 88 | tumor | Pre | resistant | NA |
| x |  |  |  | 85 | tumor | Post | resistant | 47 |
| x |  |  |  | 63 | tumor | Pre | resistant | NA |
| x |  |  |  | 76 | tumor | Post | resistant | 144 |
| x |  |  |  | 54 | tumor | Pre | resistant | NA |
| x |  |  |  | 54 | tumor | Post | resistant | 2035 |
| x |  |  |  | 61 | tumor | Post | resistant | 173 |
| x |  |  |  | 70 | tumor | Pre | resistant | NA |
| x |  |  |  | 72 | tumor | Post | resistant | 152 |
| x |  |  |  | 46 | tumor | Pre | resistant | NA |
| x |  |  |  | 47 | tumor | Post | resistant | 134 |
| x |  |  |  | 61 | tumor | Post | resistant | 95 |
| x |  |  |  | 63 | tumor | Pre | resistant | NA |
| x |  |  |  | 61 | tumor | Post | resistant | 115 |
| x |  |  |  | 65 | tumor | Post | resistant | 91 |
| x |  |  |  | 63 | tumor | Pre | resistant | NA |
| x |  |  |  | 50 | tumor | Post | resistant | 324 |
| x |  |  |  | 65 | tumor | Post | resistant | 63 |
|  | x |  |  | 73 | tumor | Post | resistant | 209 |
|  | x |  |  | 82 | tumor | Post | resistant | 159 |
|  | x |  |  | 86 | tumor | Pre | NA | NA |
|  | x |  |  | 86 | tumor | Pre | NA | NA |
|  | x |  |  | 82 | tumor | Post | resistant | 700 |
|  | x |  |  | 92 | tumor | Pre | NA | NA |
|  |  | x |  | 69 | urine | Pre | resistant | NA |
|  |  | x |  | 69 | urine | Post | resistant | 40 |
|  |  | x |  | 83 | urine | Pre | sensitive | NA |
|  |  | x |  | 83 | urine | Dose 6 | sensitive | NA |
|  |  | x |  | 83 | urine | Post | sensitive | 40 |
|  |  | x |  | 51 | urine | Pre | sensitive | NA |
|  |  | x |  | 52 | urine | Dose 6 | sensitive | NA |
|  |  | x |  | 52 | urine | Post | sensitive | 41 |
|  |  | x |  | 72 | urine | Pre | sensitive | NA |
|  |  | x |  | 72 | urine | Dose 6 | sensitive | NA |
|  |  | x |  | 72 | urine | Post | sensitive | 43 |
|  |  | x |  | 72 | urine | Post | sensitive | 43 |
|  |  | x |  | 79 | urine | Pre | sensitive | NA |
|  |  | x |  | 81 | urine | Pre | sensitive | NA |
|  |  | x |  | 82 | urine | Pre | sensitive | NA |
|  |  | x |  | 82 | urine | Dose 6 | sensitive | NA |
|  |  | x |  | 71 | urine | Pre | sensitive | NA |
|  |  | x |  | 71 | urine | Dose 6 | sensitive | NA |
|  |  | x |  | 71 | urine | Post | sensitive | 61 |
|  |  | x |  | 62 | urine | Pre | sensitive | NA |
|  |  | x |  | 74 | urine | Dose 6 | sensitive | NA |
|  |  | x |  | 74 | urine | Pre | sensitive | NA |
|  |  | x |  | 74 | urine | During (dose 6) | sensitive | NA |
|  |  | x |  | 72 | urine | Dose 1 | sensitive | NA |
|  |  | x |  | 72 | urine | Dose 6 | sensitive | NA |
|  |  | x |  | 72 | urine | Post | sensitive | 35 |
|  |  | x |  | 59 | urine | Pre | sensitive | NA |
|  |  | x |  | 69 | urine | Pre | sensitive | NA |
|  |  | x |  | 69 | urine | Dose 6 | sensitive | NA |
|  |  | x |  | 90 | urine | Pre | sensitive | NA |
|  |  |  | x | 83 | urine | Post | resistant | unknown |
|  |  |  | x | 82 | urine/tumor | Post | resistant | 398 |
|  |  |  | x | 80 | tumor | Post | resistant | 1341 |
|  |  |  | x | 74 | tumor | Post | resistant | 86 |
|  |  |  | x | 90 | tumor | Post | resistant | 1303 |
|  |  |  | x | 74 | urine | Post | resistant | 147 |
|  |  |  | x | 90 | urine/tumor | Post | resistant | 1364 |

**Table S2:**
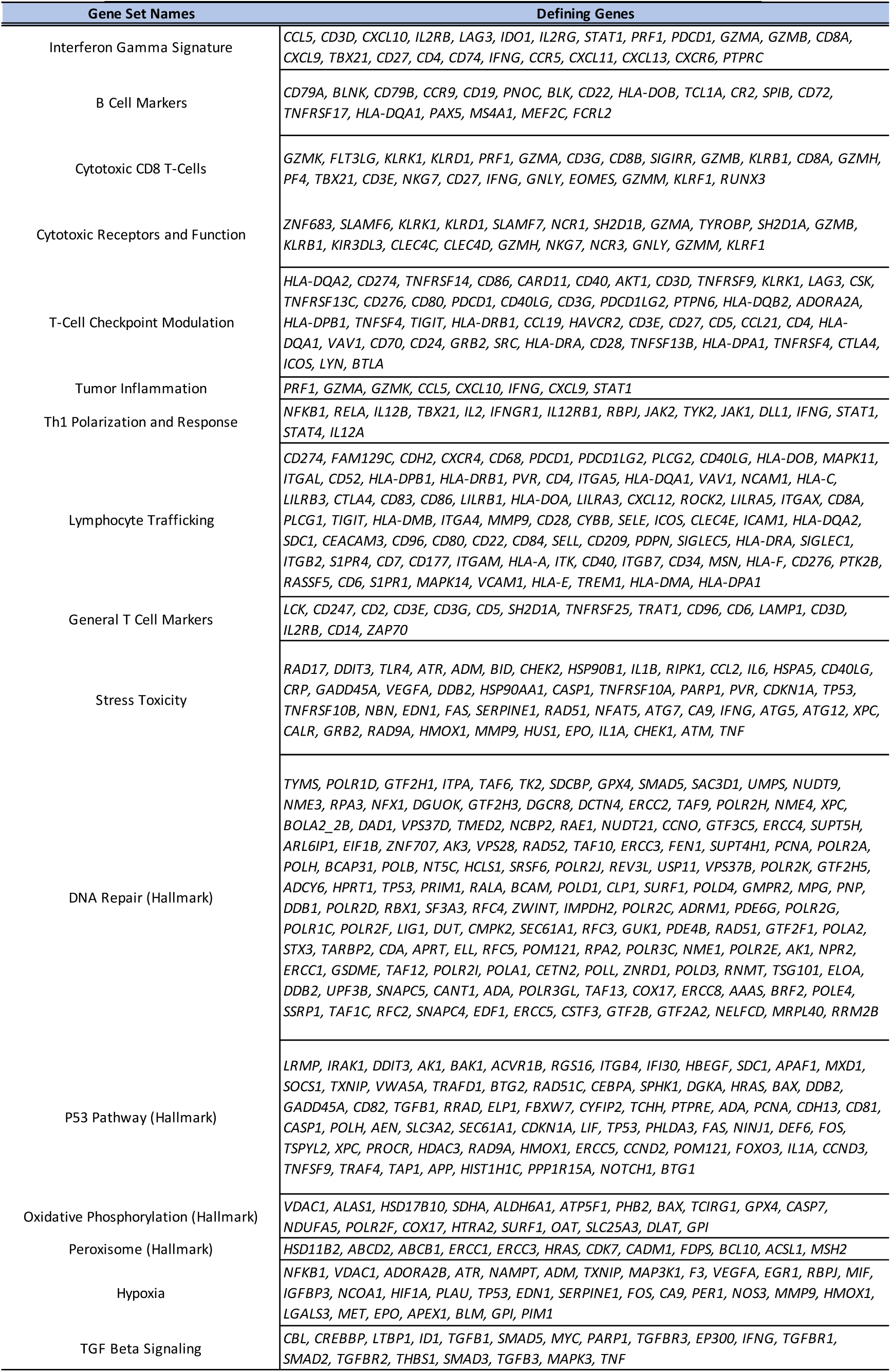
Genes defining gene-set enrichment pathways.

